# High-Resolution Transcriptional Impact of AIRE: Effects of Pathogenic Variants p.Arg257Ter, p.Cys311Tyr, and Polygenic Risk Variant p.Arg471Cys

**DOI:** 10.1101/2024.12.09.627575

**Authors:** Amund Holte Berger, Bergithe Eikeland Oftedal, Anette Susanne Bøe Wolff, Eystein Sverre Husebye, Per Morten Knappskog, Eirik Bratland, Stefan Johansson

## Abstract

The Autoimmune Regulator, AIRE, acts as a transcriptional regulator in the thymus, facilitating ectopic expression of thousands of genes important for the process of negative T-cell selection and immunological tolerance to self. Pathogenic variants in the gene encoding AIRE are causing Autoimmune polyendocrine syndrome type 1 (APS-1), defined by multiorgan autoimmunity and chronic mucocutaneous candidiasis. More recently, Genome Wide Association Studies (GWAS) have also implicated AIRE in several common organ-specific autoimmune diseases including Autoimmune primary adrenal insufficiency, type 1 diabetes and pernicious anemia. We developed a highly sensitive cell-system approach based on HEK293FT cells transfected with *AIRE* that allowed us to characterise and functionally evaluate the transcriptional potential of genetic variants in the *AIRE* gene. We confirm that our cell system recapitulates the expression of the vast majority of known AIRE induced genes including well-characterised tissue restricted antigens (TRAs), but also increases the total number of identified AIRE induced genes by an order of magnitude compared to previously published strategies. The approach differentiates between categories of *AIRE* variants on the transcriptional level, including the nonsense variant p.R257* (near complete loss of function), the p.C311Y variant associated with dominantly inherited APS-1 (severely impaired function), and the polygenic risk variant p.R471C (slightly increased function) linked to common organ-specific autoimmunity. The increased activity of p.R471C compared to wildtype indicates different molecular mechanisms for monogenic and polygenic AIRE related autoimmunity.

## Introduction

Epithelial cells of the thymus stroma, specifically medullary thymic epithelial cells (mTECs), are instrumental for the generation of a repertoire of self-tolerant T-cells [1]. mTECs are able to turn on the expression of thousands of genes normally restricted to cell lineages of specialized tissues, collectively referred to as tissue restricted antigens (TRAs) [2]. The precise molecular events that drive this transcriptional plasticity in mTECs are incompletely understood, although studies of the rare monogenic condition Autoimmune polyendocrine syndrome type 1 (APS-1) have identified a crucial role for the AutoImmune Regulator (AIRE) in these processes. APS-1, also known as autoimmune polyendocrinopathy candidiasis-ectodermal dystrophy (APECED, OMIM 240300), is clinically characterized by a wide range of autoimmune manifestations, affecting both endocrine and non-endocrine tissues [3].

APS-1 can be inherited in both autosomal dominant and recessive manners depending on the molecular effect of the pathogenic genetic variants in AIRE, encoding the AutoImmune Regulator (AIRE) protein [4, 5, 6, 7, 8]. In addition to the monogenic disorder APS-1, genetic variations in AIRE have recently been associated with more common organ-specific autoimmune diseases such as Addison’s disease, pernicious anaemia and type 1 diabetes [9, 10, 11]. In particular the p.R471C variant, with a frequency of 1-2% in Northern European populations, have been associated with more than 3-fold increased risk of Addisońs disease, almost 2-fold increased risk of pernicious anaemia and 1.5-fold increased risk of type 1 diabetes. Thus, the extensive heterogeneity in effects, ranging from autosomal dominant to bi-allelic and polygenic risk-variants within AIRE has created a need for functional characterization and clinical classification of AIRE variants.

AIRE is predominantly expressed in the mTECs where it facilitates the expression of TRAs normally restricted to alternative cell lineages representing most tissues and organ systems [12]. AIREs amino acid sequence and predicted domain structure are indicative of a chromatin associated factor, and since its discovery, it has been assumed that AIRE is involved in transcriptional regulation [4, 13]. However, AIRE’s interaction with DNA appears to be sequence independent (at least in vitro [14]), and the transcriptional footprint of AIRE is far too broad for a classical transcription factor [12, 15]. AIRE contains a so-called SAND (Sp100, AIRE, NucP41/P75, DEAF1) domain, which is found in a number of nuclear proteins thought to play important roles in chromatin-dependent transcriptional regulation [16]. Although, AIRE’s SAND domain lacks the conserved DNA binding motif consisting of the amino acids Lys-Trp-Asp-Lys, suggesting a different role than sequence dependent DNA binding. Additionally, AIRE contains two plant homeodomain (PHD) zinc finger domains, PHD1 and PHD2, indicating a potential to interact with chromatin through the binding of histone tails [4]. However, only PHD1 has convincingly been shown to have histone tail-binding activity as it preferentially binds to the unmodified N-terminal tail of H3, a hallmark of silent chromatin [14]. PHD2 has instead been suggested to act as an adaptor for other multiprotein complexes involved in transcriptional elongation [17]. Finally, AIRE also contains a caspase activation recruitment domain (CARD), which is required for AIRE homo-multimer formation and transcriptional activity [18]. Taken together, and further corroborated by a number of state-of-the-art single cell analysis experiments, it seems that AIRE is indirectly inducing TRA expression by ensuring transcriptional heterogeneity of mTECs [2, 19, 20]. This is achieved by the repurposing of general transcriptional mechanisms and by alteration of the chromatin landscape, such as pause-release of stalled RNA polymerase II and enhancer-promoter looping [14, 21, 22, 23]. Recently, it was also discovered that AIRE is preferentially recruited to its target genes by promoters already poised for transcription (independently of AIRE, characterized by Z-DNA enrichment and double-stranded breaks in DNA [24]. Still, in spite of numerous ingenious studies based on mTECs from genetically modified mouse models that has contributed immensely to the knowledge of functional aspects of AIRE, its exact way of action is incompletely understood at several levels. In particular the definition of bona fide AIRE targets is problematic due to the low and promiscuous expression levels of TRAs in the mTECs.

Transcriptomics is particularly well-suited for the illumination of AIRE’s molecular properties, given the vast number of induced or perturbed targets in the presence of AIRE in a given cell. However, in order to avoid the sheer complexity in the interpretation of transcriptomics data derived from a highly heterogeneous cellular population such as mouse mTECs, additional complementary cell systems are needed for AIRE-related studies. In fact, cell-line based models may in some ways be better suited for the exploration of the molecular mechanism of AIRE, as they are not limited by the number of AIRE expressing cells in a given tissue (such as mTECs) and can be easily genetically altered. Therefore, we have designed a simple and accessible cell system exploiting the homogeneous nature of the widely used and well characterized cell-line HEK293, and a plasmid conferring high levels of AIRE without leading to overloading of protein synthesis. We have chosen to study three different AIRE variants in addition to the WT, representing different proposed molecular mechanisms and different associated clinical phenotypes: the nonsense variant p.R257* located within the CARD domain [25], the dominant negative variant p.C311Y located within the PHD1 domain [26], and the polygenic risk variant p.R471C located within the PHD2 domain [17].

In the context of the functional characterization of AIRE, this offers a simple solution to the problems represented by the considerable heterogeneity of mTECs and their highly diverse and complex spectrum of TRA expression and transcriptional networks, as well as different maturation stages and subpopulations. We show that our AIRE transfected HEK293FT cell line system can be used to give increased insight to the molecular function of AIRE and to differentiate between AIRE variants based on their transcriptomic properties.

## Methods

### Plasmid amplification and mutagenesis

pSF-UB plasmids were purchased from OXGENE with and without *AIRE* cDNA inserts. Empty pSF-UB plasmids were used as negative control, while pSF-UB plasmids containing *AIRE* were used for all *AIRE* WT transfections and mutagenesis to create the AIRE mutations p.R257*, p.C311Y and p.R471C. Mutagenesis was performed using QuikChange II Site Directed Mutagenesis kit from Agilent Technologies. Plasmids were amplified using TOP10 competent E. coli cells from Thermo Fisher and purified using a QIAprep Spin Miniprep Kit from Qiagen. The sequence of *AIRE* and mutated *AIRE* containing plasmids were confirmed by Sanger sequencing.

### Cell culture and transfection

HEK293FT human embryonic kidney cells (RRID: CVCL_6911) were grown in a medium consisting of Dulbecco’s Modified Eagle Medium supplemented with 4.5 g/l D-Glucose, Pyruvate, 10% Fetal Bovine Serum (FBS), and 1% Penicillin-Streptomycin. The cells were incubated at 37 °C in a 5% CO2 humidified incubator until reaching 80-100% confluency. The cells were subsequently counted using a Scepter cell counter from Merck and transferred to 6-well plates, where each well was seeded with 6×105 cells. After 24h, the cells were transfected using the Lipofectamine 2000 Transfection Reagent from Thermo Fisher with 2.5 *µ*g DNA and 12 *µ*l Lipofectamine.

### Western immunoblotting

Transfected cells grown for 48 hours were removed from the growing surface by flushing, then washed using centrifugation and resuspension in PBS (Cat#: D8537, Merck). Cells were lysed using cOmplete lysis buffer (Cat#: 04719956001, Merck) for 30 minutes on ice. LDS-PAGE was performed using NuPAGE MOPS SDS Buffer Kit (Cat#: NP0050) according to its instructions and run on a NuPage 10% Bis-Tris gel (Cat#: NP0301BOX, Thermo Fisher) with SeeBlue Plus2 Pre-Stained Protein Standard (Cat#: LC5925, Thermo Fisher) at 180V for 1 hour and 10 minutes. Samples were transferred to a PVDF membrane using the iBlot dry blotting system (Cat#: IB401002, Thermo Fisher). Washing steps were performed using Pierce TBS Tween 20 (Cat#: 28360, Thermo Fisher) while blocking was performed using SuperBlock Blocking Buffer (Cat#: 37517, Thermo Fisher) for 1 hour on a shaker. Membranes were incubated with primary antibody in TBS-T with 5% BSA overnight on a shaker in a cold room. The primary antibodies used were mouse *α*-GAPDH (Cat#: MAB374, Merck) in a 1:1000 dilution, goat *α*-AIRE (Cat#: MAB155, Merck) in a 1:500 dilution. The membranes were rewashed, then incubated with the secondary antibodies in TBS-T with 5% BSA for 1 hour on a plate shaker. The secondary antibody used was goat *α*-Mouse antibody (Cat#: 626520, Thermo Fisher) conjugated with Horse Radish Peroxidase (HRP) in a 1:2000 dilution. After rewashing the membranes were soaked in Pierce ECL Western Blotting Substrate (Cat#: 32106, Thermo Fisher) and imaged using a Bio-Rad Chemidoc MP.

### RNA isolation

Transfected cells grown for 48h were harvested by flushing, then transferred to 15ml tubes where they were centrifuged at 300g for 7 minutes before being resuspended in cold PBS. RNA was isolated using Qiagen RNeasy Mini Kit. A Qiagen QIAshredder spin column was used to homogenise the samples, and a Qiagen RNase free DNase solution was used in order to digest any genomic DNA. The NanoDrop microvolume spectrophotometer (Thermo Fisher) was used to check the RNA quantitation. In order to ensure that high-quality RNA was used, the samples were also analysed using an Agilent Bioanalyzer 2100 with an Agilent RNA 6000 Nano kit.

### Real-time quantitative PCR

RNA from an independent experiment was analysed using qPCR consisting of a two-step process using Thermo Fisher Superscript VI VILO cDNA Synthesis Kit (Cat#: 11754050) followed by Thermo Fisher TaqMan Universal PCR Master Mix (Cat#: 4304437) and a variety of TaqMan probes according to product info sheet and user guide. qPCR was performed using Applied Biosystems QuantStudio 5 and the Thermo Fisher Cloud software. Three technical controls for each condition were used, in addition to no template control (NTC) and no reverse transcriptase control (-RT). TaqMan probes consisted of probes for the genes *AIRE* (Cat#: Hs00230829_m1), *CCNH* (Cat#: Hs00236923_m1), *S100A8* (Cat#: Hs00374264_g1), *IGFL1* (Cat#: Hs01651089_g1), *KRT14* (Cat#: Hs00265033_m1), and *GAPDH* (Cat#: Hs99999905_m1). The qPCR results were processed using the ΔΔCt method in R, and normalised against the *GAPDH* housekeeping gene, while fold change was calculated between *AIRE* transfected and empty plasmid transfected control. Data was prepared in R [27] (v.4.0.2) and Rstudio [28] (v.1.4.1103) using the tidyverse [29] (v. 1.3.1) package, then plotted using ggplot2 [30] (v.3.3.5) with the additional packages RColorBrewer [31] (v. 1.1-2) and scales [32] (v. 1.1.1).

### RNAseq

Illumina TruSeq Stranded mRNA Library Prep kit together with Illumina TruSeq RNA CD Index Plate index adapters was used for library preparation. Sequencing was performed with 12 biological replicates using an Illumina HiSeq 4000 sequencer, with a read depth of approximately 100 million reads per sample.

### RNA sequencing alignment

Quality control of fastq files were performed using FastQC [33] (v.0.11.9)and MultiQC [34] (v.1.12). Fastq files were subsequently aligned using the Kallisto [35] pseudoaligner (v.0.46.2) to the GRCh38.p12 (release 100) reference genome provided by ensembl [36]. All subsequent analysis were performed using R [27] (v.4.0.2) and Rstudio [28] (v.1.4.1103), where tximport [37] (v.1.18) was used to import Kallisto output files into R and aggregate transcript data to the gene level using the “hsapiens_gene_ensembl” gene annotation data accessed from biomaRt [38] (v.2.46.3).

### Differential expression analysis

Differential expression analysis was performed using DESeq2 [39] (v.1.30.1) with the design formula “~mutation” with *AIRE* WT and variant genotypes separately vs empty control and *AIRE* variants separately vs *AIRE* WT samples. Significance testing was performed using a Wald test with an FDR threshold of 5%. Log2foldchange shrinkage was performed using apeglm [40] (v.1.12.0). Result tables were generated using tidy data principles with the tidyverse [29] package(v.1.3.1), and volcano, hexbin, violin and dotplots were generated using ggplot2 [30] (v.3.3.5). Colour scales used were generated using the packages RColorBrewer [31] (v.1.1-2) and Viridis [41] (v.0.6.2). Plots were also generated with the help of the additional packages ggrepel [42] (v.0.9.1), cowplot [43] (v.1.1.1), ggtext [44] (v.0.1.1), and scales [32] (v.1.1.1). Venn diagrams were generated using the VennDiagram [45] (v.1.7.0) package, but plotted in ggplot using draw_grob from cowplot.

### Categorisation of genes according to tissue specificity and restriction

As AIRE induced gene expression has been considered to predominantly consist of tissue restricted antigens (TRA), i.e. genes only expressed in a small subset of tissues, we classified genes as tissue enriched [46], group enriched [47], or tissue enhanced [48] according to their entries by the Human Protein Atlas (HPA) [49, 50]. Tissue enriched genes were defined as genes with a fourfold increase in mRNA expression in one tissue compared to any other tissue, group enriched genes as genes with a four-fold increase in mRNA expression in 2–5 tissues compared to all other tissues, and tissue enhanced genes as genes with a four-fold increase in mRNA in one tissue compared to the average level in all other tissues.

## Results

### Expression of functional AIRE in transiently transfected HEK293FT cells

To induce functional AIRE expression, HEK293FT cells were transfected with a pSF-UB plasmid containing *AIRE* cDNA. In addition to *AIRE* wildtype (WT), we also included the variants p.R257*, p.C311Y and p.R471C, to investigate whether transient transfection could be used to differentiate between WT and variants. The nonsense variant p.257* was selected as a representative of recessive variants with a large effect, p.C311Y was selected as a dominant variant with a moderate effect and p.R471C was selected as a common variant with small effect. Western blot analyses using a polyclonal anti-AIRE antibody targeting the N-terminal of human AIRE and protein lysates from cells transfected with pSF-UB plasmids encoding wild-type (WT) AIRE and AIRE variants revealed bands corresponding to the expected molecular weight for all AIRE variants (**Fig. 1a**).

**Figure 1:**
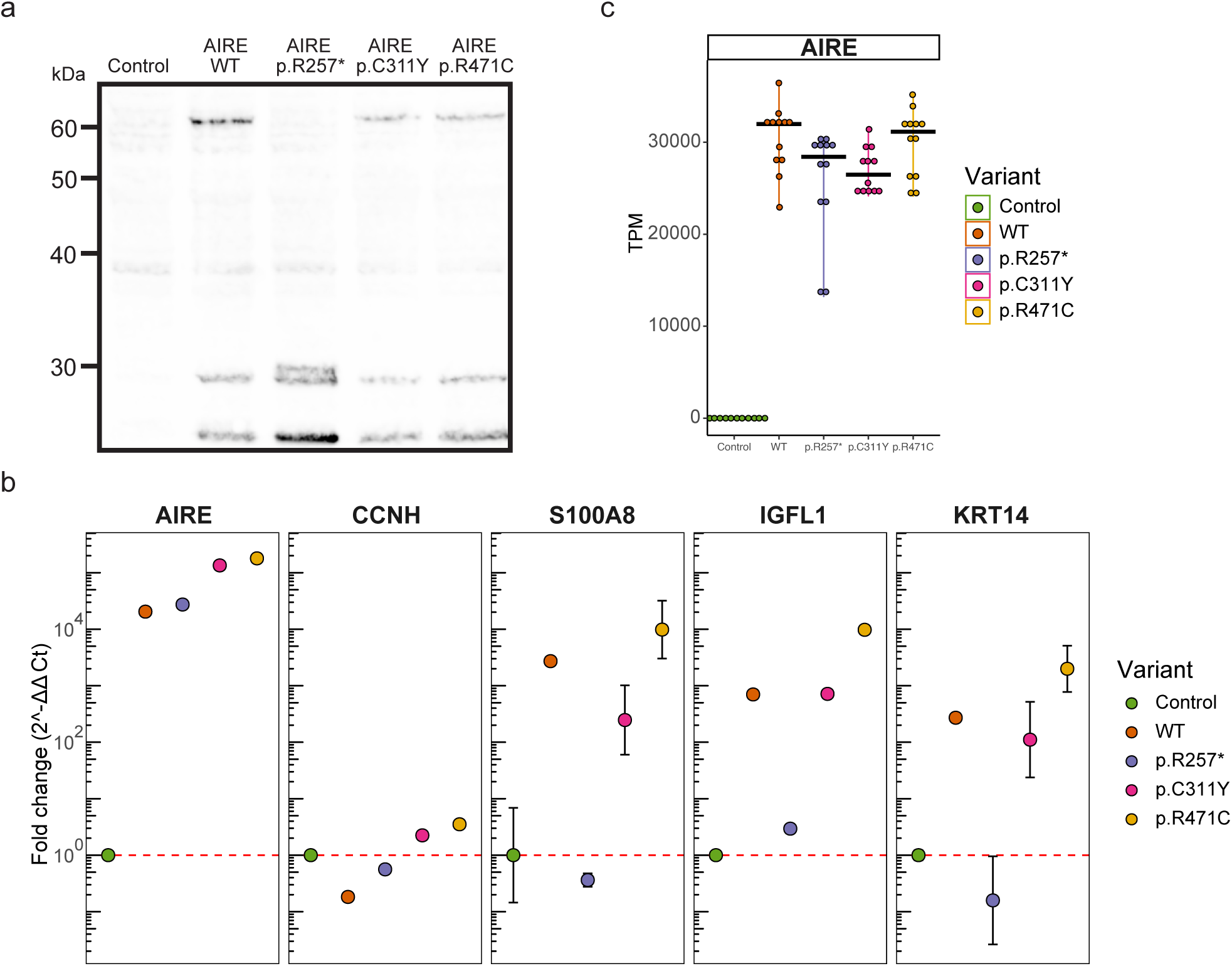
Expression of functional AIRE. **a** Western blot of cellular lysates containing AIRE wildtype and variants p.257*, p.C311Y, and p.R471C from transfected HEK293FT cells. AIRE wildtype, p.C311Y and p.R471C exhibit full length bands at around 60 kDa, while the truncated p.R257* exhibits a band at approximately 30kDa. **b** qPCR of RNA from transfected HEK293FT cells. *CCNH* used as negative (i.e. non-AIRE induced) control, *S100A8*, *IGFL1* and *KRT14* used as known (i.e. canonical) AIRE regulated genes. **c** Expression of *AIRE* in transcripts per million (TPM) as determined by an independent RNAseq experiment of RNA from transfected HEK293FT cells.

Quantitative PCR on RNA from cells transfected with each of the four variants showed that AIRE expression was increased by 10^3^-10^5^ fold for all *AIRE* variants compared to empty vector control (**Fig. 1b**). For the canonical AIRE induced genes *S100A8*, *IGFL1* and *KRT14*, expression increased dramatically in WT, p.C311Y and p.R471C transfected cells, while no induction of gene expression could be seen in the cells with the nonsense mutation p.R257*. For *S100A8* and *KRT14*, but not *IGFL1*, induction was greatly reduced in p.C311Y transfected cells compared to WT. Notably, cells transfected with *AIRE* variant p.R471C exhibited consistently higher expression of AIRE induced genes than WT. An independent RNAseq experiment replicated the robust expression of the *AIRE* variants in transfected cells (**Fig. 1c**). Furthermore, RNAseq confirmed expression of a range of verified and well-characterized molecular partners in AIRE-mediated transcriptional regulation, further supporting the suitability of the cell-system for biochemical and molecular studies of AIRE (**Supplementary Figure A1**). For most of the molecular partners of AIRE, the overexpression of *AIRE* did not influence gene expression levels. The exception was tissue enhanced transcription factors such as *HNF1A*, *HNF1B* and *HNF4A*, that were all clearly induced by AIRE in our system.

### AIRE variants induce a spectrum of variant-dependent transcriptional effects, reflecting the associated clinical phenotypes

After observing robust expression of functional AIRE in transfected HEK293FT cells, we wanted to identify the global expression profiles induced by different *AIRE* variants. RNA sequencing was performed on high quality RNA derived from HEK293FT cells transfected with empty control pSF-UB plasmid, and pSF-UB plasmids containing cDNA for *AIRE* WT, p.R257*, p.C311Y, or p.R471C (**Fig. 1c**). To enable the robust discovery of low abundance transcripts, typical for AIRE induced genes, a total of 12 replicates per variant were included at a read depth of approximately 100 million.

Principal component analysis (PCA) revealed a clear separation between the WT and variant transfected cells compared to empty control using PC1 and PC2 (**Fig. 2a**). The overall expression pattern induced by the nonsense mutation p.R257* indicates that it clusters more closely to the empty control, the missense mutation p.C311Y displays intermediate clustering between p.R257* and WT, while WT and p.R471C induced similar expression patterns.

**Figure 2:**
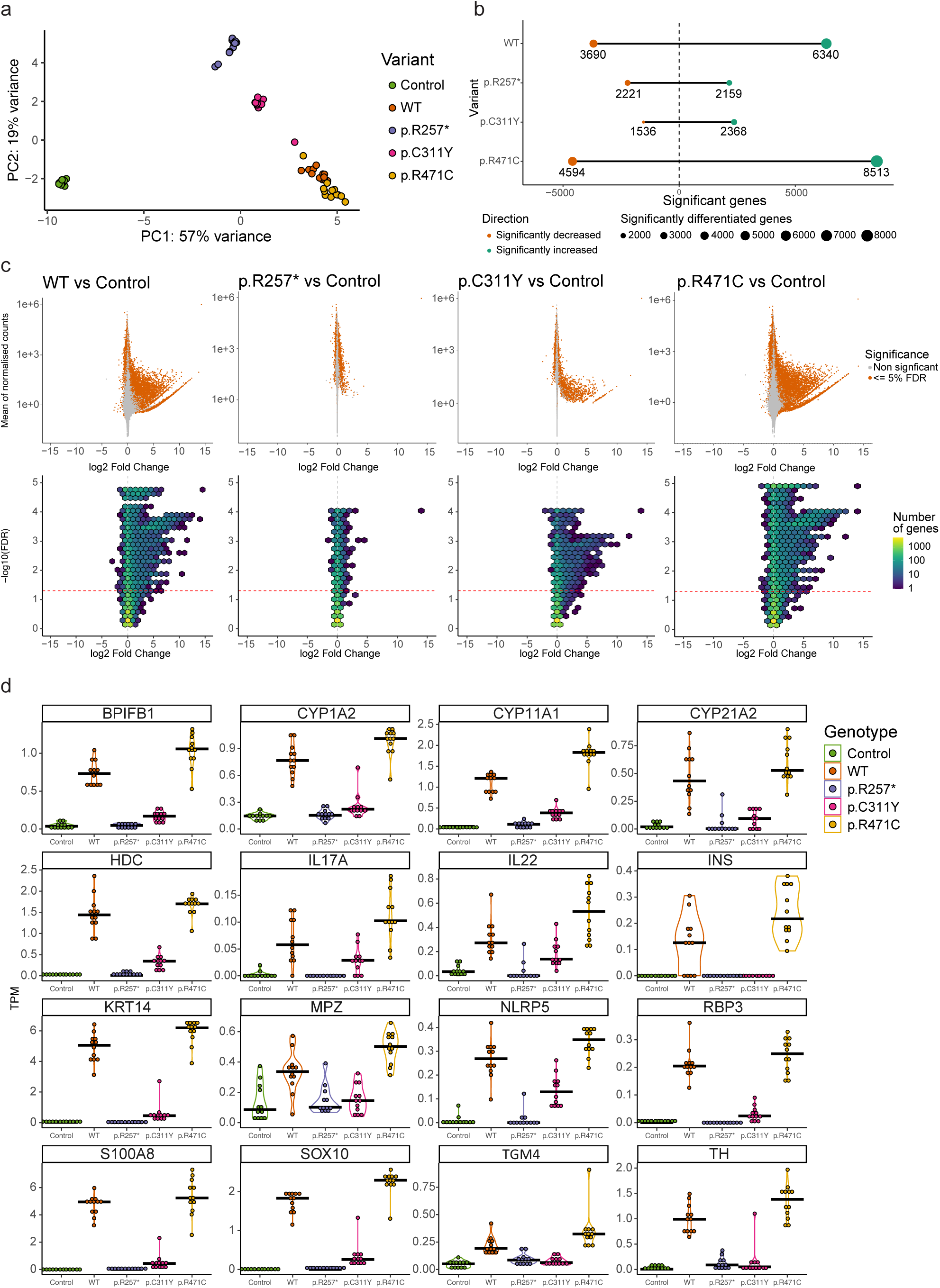
AIRE induced expression and transcriptional effects of AIRE variants revealed by RNA sequencing. **a** PCA plot of HEK293FT samples transfected with empty control plasmid, AIRE wildtype or AIRE variants p.R257*, p.C311Y, p.R471C (n = 12 for all conditions, *AIRE* levels seen in **Fig. 1c**) **b** Number of significant differentially expressed genes of *AIRE* WT and *AIRE* variant transfected cells compared to control cells (FDR <= 0.05). **c** MA plot (upper row) and Hex bin Volcano plots (lower row) displaying differentially expressed genes induced by AIRE wildtype and variants compared to empty vector control. **d** Differential expression of a selected subset of AIRE-relevant genes (encoding known autoantigens in APS-1 and/or previously reported to be AIRE-regulated) by different AIRE variants, in transcripts per million (TPM).

Differential gene expression analysis revealed a massive upregulation of genes induced by WT AIRE compared to empty vector. A total of 6340 genes were significantly higher expressed and 3690 lower expressed at a 5% FDR level (**Fig. 2b**), with up to 15 times log2 fold change (Log2FC) increased expression levels for certain genes (**Fig. 2c**). The non-sense mutation p.R257* however, exhibited a different pattern characterized by over-all low log2 fold change and symmetrical effect in directions of expressed genes (2159 vs 2221 respectively) compared to empty vector (**Fig. 2b-c**). The missense mutation p.C311Y induced significantly increased expression levels of 2368 genes accompanied by significantly reduced expression of 1536 genes, both categories with log2 fold changes similar to those induced by AIRE WT (**Fig. b-c**). The p.R471C variant exhibited a pattern distinct from the two pathogenic variants, with the induction of significantly higher expression of 8513 genes and significantly lower expression of 4594 genes compared to empty vector control, with log2 fold changes similar to, or higher, than those induced by WT.

**Figure 2d** shows the absolute expression data in transcript per million (TPM) of a curated list of genes previously shown to be induced by AIRE in both human and murine systems and genes encoding known APS-1 autoantigens representing a wide range of tissues such as adrenal cortex, ovaries, lung, liver, pancreas, and parathyroid. While AIRE WT clearly induced or enhanced the expression of all selected genes, the p.R257* variant induced no or little changes in expression levels. The p.C311Y missense mutation displayed limited ability to induce or enhance gene expression, and was unable to induce any expression of some of the selected AIRE targets above the levels of empty vector control. p.R471C, comparatively, consistently induced or enhanced gene expression greater than AIRE WT.

### The developed cell system recapitulates findings from earlier studies, but expands the range of AIRE induced target genes

To analyse the performance of our AIRE induced expression cell system we compared our data set to data from a published comparable cell system Guha et al. (2017) [51]. In their study, HEK293 cells were transfected with *AIRE* WT and compared to untransfected cells. The main focus of the study by Guha and colleagues was the interaction of AIRE with topoisomerase2 (TOP2) in order to induce double-stranded breaks leading to enhanced chromatin access. Treating *AIRE* transfected cells with the TOP2 inhibitor etoposide was found to increase AIRE induced gene expression. To avoid systematic differences due to different methodology, we reanalysed their raw data using our own differential expression analysis pipeline and only used data from cells not treated with etoposide. This re-analysis of their data identified 507 significantly up-regulated genes and 5 down-regulated genes at an FDR of 5% (**Fig. 3a**). These numbers are slightly lower but comparable to what was described in the original publication (n=691 induced genes with an FDR of 10%). A comparison with our study reveals that the vast majority (88%) of AIRE WT induced genes in Guha et al. 2017 were also identified in our cell system (WT vs control) (**Fig. 3b**). This confirms that the transfection-based approach developed by us recapitulates findings of earlier studies yet offers an additional layer of high-resolution data.

**Figure 3:**
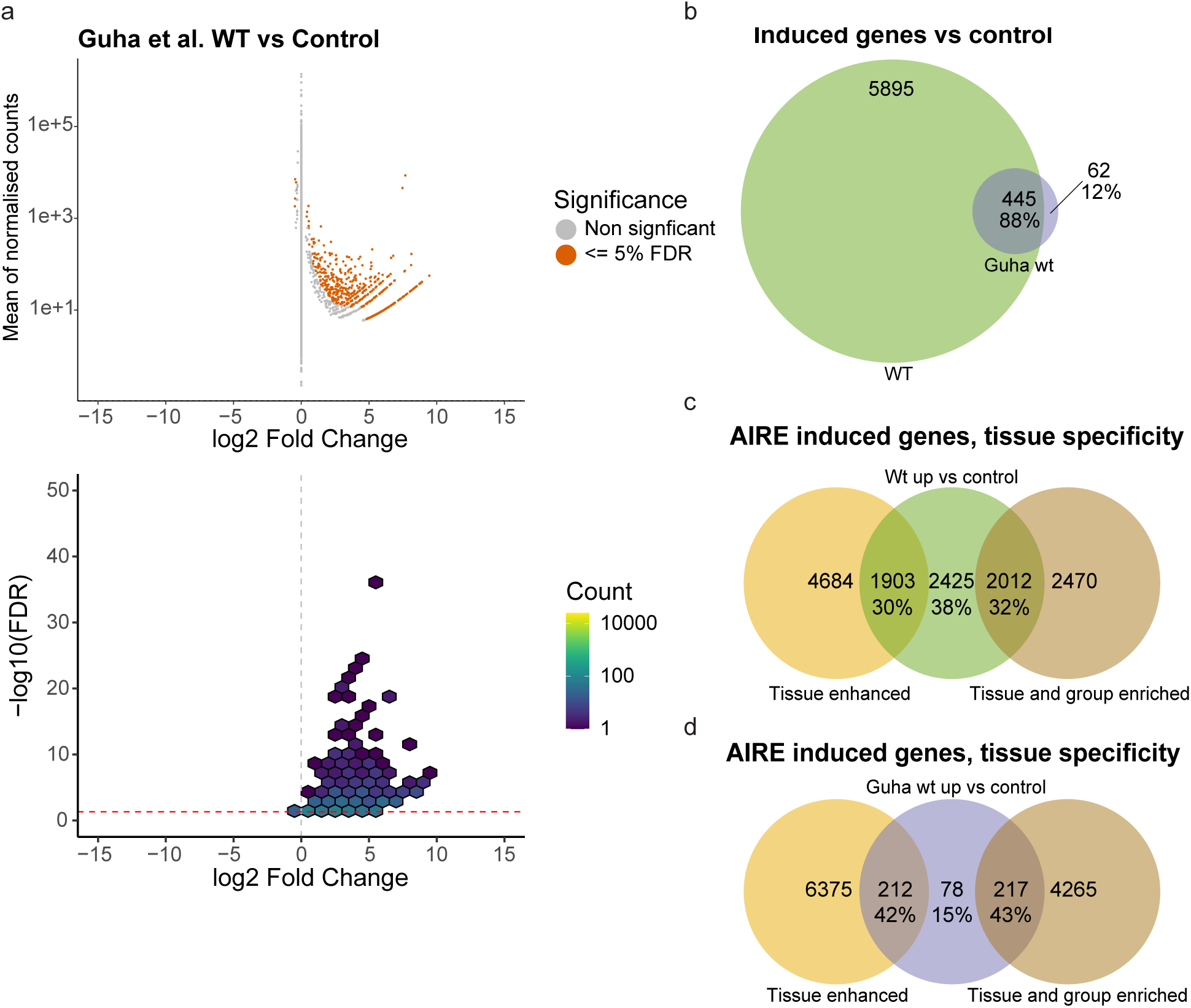
Comparisons with previous AIRE cell studies, and proportion of tissue restricted genes. **a** RNAseq from raw data provided by Guha et al. (2017) [51]. **b** Overlap of genes found significantly expressed in cells transfected with wildtype *AIRE* compared to control in this study and in Guha et al. **c** Genes significantly expressed in cells transfected with wildtype *AIRE* compared to cells transfected with empty vector control, divided into genes classified as tissue enhanced, or tissue and group specific genes according to the Human Protein Atlas. **d** Genes induced by AIRE WT in Guha et al. divided into genes classified as tissue enhanced, or tissue and group specific genes according to the HPA.

AIRE induced gene expression has been considered to be predominantly confined to Tissue Restricted Antigens (TRA), i.e. genes normally only expressed in a small subset of tissues. To explore this view, we compared the 6340 AIRE induced genes (defined as significantly higher expressed in AIRE WT transfected cells) to proteins classified as tissue enriched, group enriched, or tissue enhanced by the Human Protein Atlas (HPA). Thirty percent of genes induced by AIRE WT were classified as tissue enhanced by HPA, while 32% were classified as either tissue enriched or group enriched, the remaining 38% were not classified as either (**Fig. 3c**) compared to 27%, 8% and 65% respectively for all genes expressed at levels greater than 1 TPM in empty control vector transfected cells (data not shown).

As a comparison, we analysed the distribution of AIRE WT induced genes in the data set reported by Guha et al. (using our analysis pipeline). Here, a greater proportion of significantly upregulated genes were classified as tissue enhanced (42%) and tissue or group enriched (43%) while only 15% were outside these groups (**Fig. 3d**). The lower proportion of non-tissue restricted genes in Guha et al. compared to the 38% found in our cell-assay might be explained by the higher resolution of our dataset, providing sufficient statistical power to reveal that AIRE not only is able to induce so-called ectopic expression of tissue-specific genes, but also to induce and/or enhance expression of genes not considered tissue-specific.

### AIRE is capable of inducing low level transcription of non-tissue restricted genes

While the classical fold-change measure emphasizes the relative differences between induced genes, the absolute expression in transcripts per million (TPM) allows for a more direct comparison of the natural expression levels of genes induced by AIRE. The absolute expressions are depicted in **Figure 4** and shows that AIRE induces low-level increase in gene expression for the majority of targeted genes, with most genes exhibiting an increase below 1 TPM and a median increase of 0.39 TPM (**Fig. 4a-c**) from a median baseline TPM in control of 0.11 to a median TPM of 0.51 in WT. However, there are some exceptions where absolute gene expression is increased by as much as 57 TPM. Comparatively, genes significantly induced by p.C311Y had a median baseline of 0.12 TPM in control and was increased by a median 0.24 TPM to a median level of 0.36 TPM, while significant genes induced by p.R471C started at a median baseline of 0.16 TPM, increased by a median 0.46 TPM to a median level of 0.62 TPM (**Fig. 4a & c**). To further investigate the differences between individual genes classified into the tissue-specific categories, the differential expression data of individual genes in *AIRE* WT transfected cells and empty vector control HEK293FT cells were further divided into the three groups defined above (**Fig. 4d**). Among these groups, tissue and group enriched genes had the lowest baseline expression with a median TPM of 0.04 in empty vector control cells and a median increase of 0.29 TPM in WT transfected cells to a median level of 0.33 TPM, consistent with their classification as tissue restricted. Tissue enhanced genes had a baseline median expression of 0.14 TPM with a median AIRE induced increase of 0.40 TPM to a median level of 0.53 TPM, while the genes not classified as tissue enriched or enhanced had a baseline median expression level of 0.65 TPM and a median increase of 0.62 TPM to a median level of 1.27 TPM. These results suggest that AIRE may have a magnifying effect not only restricted to the tissue restricted genes, but also on genes already expressed at low to moderate levels in HEK293FT cells. The low absolute induction may explain why earlier analyses are biased towards changes in tissue restricted genes where the fold-change is comparatively higher and thus power is higher.

**Figure 4:**
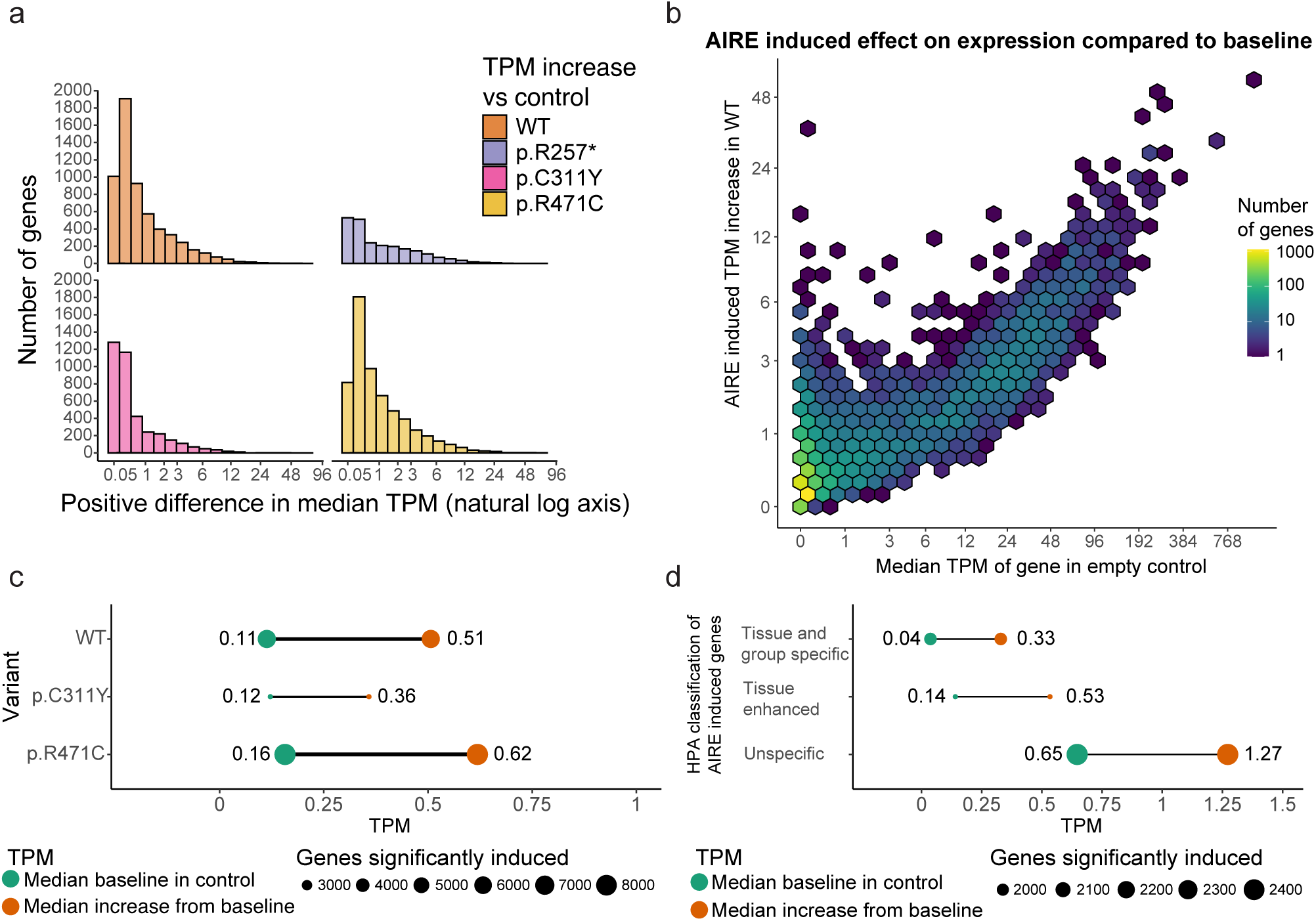
The effect of AIRE on individual genes varies greatly. **a** The median increase in absolute expression of individual genes as a result of *AIRE* WT or *AIRE* variant transfection. **b** The median increase in TPM of genes under the influence of AIRE wildtype compared to the baseline expression of those genes in empty vector control, the colour scale denotes the density of genes within the area of a hex. **c** The median level in TPM of genes significantly induced by AIRE WT and AIRE variants in control (green) and the median level in the respective AIRE variant (orange), p.R257* not shown as it exhibited a much higher baseline level. **d** TPM levels of AIRE WT induced genes separated into different levels of tissue restricted by the Human Protein Atlas, median TPM levels in control (green) and the increased median TPM levels in WT (orange)

**Supplementary Figure A1:**
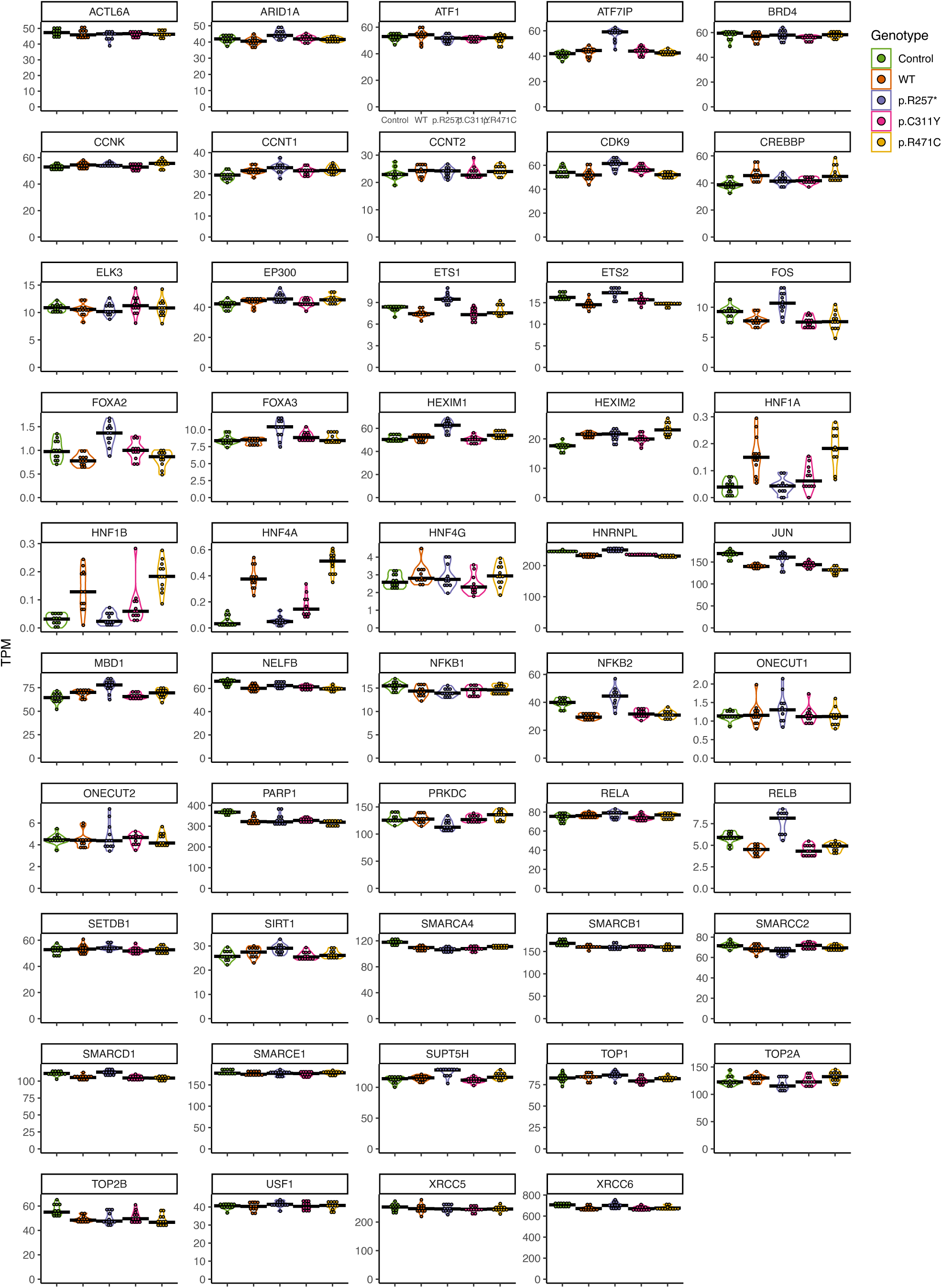
RNA expression levels of molecular partners of AIRE in HEK293FT cells. Differential expression of a selected subset of known AIRE partners by different AIRE variants, in transcripts per million (TPM)

## Discussion

The molecular mechanisms behind AIRE-mediated induction of tissue restricted antigens in the thymus are still unclear, despite more than two decades of intense research [52]. We have developed a simple, accessible and affordable cell-culture system based on transient transfection of an established human cell-line with plasmids encoding *AIRE* variants associated with different clinical phenotypes. By combining a large number of replicates and high-depth RNAseq, the developed cell assay allows for in depth analysis of differential transcriptional effects of *AIRE* variants, providing detection of both dramatic and rather subtle differences induced by the different variants. Furthermore, we present a comprehensive analysis of the transcriptional effects of AIRE at a higher level of resolution compared to earlier studies. Our results indicate that AIRE is extremely potent as a transcriptional activator across an even broader range of target genes than previously estimated [2, 51]. In addition to inducing gene expression of a multitude of tissue-restricted/tissue-enriched antigens, AIRE also seems to induce or enhance the expression of a wide range of genes that are not tissue specific. We believe this observation is in accordance with AIREs ability to act on previously poised non-coding regulatory elements and that it reflects AIREs multifaceted action on both chromatin accessibility and transcriptional initiation and elongation [53].

Numerous ingenious in vivo, in vitro and in silico experimental model systems have been described for molecular and immunological studies of AIRE [24, 12, 22, 2, 54, 55]. Together, such models have contributed to better understanding of AIRE’s function during central tolerance and beyond [52]. However, advanced model systems including those reliant on animal models might require access to considerable infrastructure not available to most standard-equipped labs. Therefore, we here propose another approach based on transient transfection of a widely used human cell-line with different variants of *AIRE*, in order to establish a manageable and reproducible assay accessible to most experimental laboratories. To this end, we utilized expression plasmids containing an ubiquitin promoter of moderate strength. Preliminary data from our laboratory (data not shown) indicated superior results from this expression vector compared to more traditional CMV promoter containing vectors, and we were able to achieve robust heterologous expression of *AIRE* in transfected HEK293FT cells, presumably without overloading the protein synthesis machinery of the host cell. *AIRE* expression was evident on both protein and mRNA levels, and AIRE activity was confirmed by assessing gene expression of canonical AIRE target genes in *AIRE* transfected cells, using both qPCR and RNAseq analyses. In addition to being permissive to *AIRE* expression, we show that HEK293FT cells also express a wide range of genes known to be critical for AIRE-mediated transcriptional regulation [52, 21]. These include components of gene repression, transcriptional elongation, mRNA splicing, chromatin structure and remodulation, and post-translational modification. A subset of these genes involved in AIRE-mediated transcriptional regulation were also shown to be induced by AIRE itself in HEK293FT cells, such as members of the hepatocyte nuclear factor (HNF-) family of transcription factors.

Importantly, we verify induction of several clinically relevant autoantigens upon *AIRE* transfection. These included genes encoding highly tissue restricted proteins such as CYP21A2/21OH (adrenal cortex) and NLRP5/NALP5 (parathyroid and ovaries), representing the main autoantigens of Addison’s disease and hypoparathyroidism, the predominant endocrinopathies of APS-1 [56, 3, 57]. As previously reported [58, 59, 60] replicates are more important than read depth for power in RNAseq experiments, so the very high sequencing depth and multiple replicates increased our power to identify much smaller alterations than previous studies and thus reveal more of the AIRE induced global expression changes. This allowed us to demonstrate that many genes encoding clinically relevant autoantigens in the context of AIRE/APS-1 were differentially induced by the different AIRE variants, reflecting their associated clinical phenotypes. To our knowledge, this is the first time AIRE has been shown to induce such a comprehensive set of clinically relevant APS-1 autoantigens. Several other well-established AIRE targeted genes, such as *KRT14* and *S100A8*, previously used as surrogate markers of AIRE activity in both mTECs and transfected cells [61, 21, 62], were also shown to be induced in our system.

While p.R257* shows little to no activity in inducing known AIRE target genes, it still presents with a large number of significantly upregulated genes compared to the control when looking at the overall differential expression analysis. These genes have a much higher average expression level in the control compared to the significantly induced genes by the other AIRE variants. As such, it is possible that these changes are the result of an artificial overabundance of a truncated protein variant (**Fig. 1a**) that consists of the CARD and partially of the SAND domain but lacks the PHD1 and PHD2 domains necessary for targeting by AIRE of inactive chromatin regions and partner interaction. It is probable that this variant would undergo nonsense-mediated decay in vivo, and as such any transcriptional effects here can be ignored. The p.C311Y variant is able to induce expression of many known AIRE targets, but to a much lower degree than AIRE WT, similar to the nearby variant p.D312A as previously described [63]. This is also consistent with its clinical presentation with a later onset and possibly milder phenotype. Unexpectedly, the common variant p.R471C consistently induces more genes and induces those genes to a larger degree than AIRE WT. As this variant is not associated with APS-1 but rather with the risk of developing common organ-specific autoimmunity [9, 10, 11] it is possible that this increased activity disrupts antigen presentation enough to increase the likelihood of escape by autoreactive T-cells in the thymus. We might speculate that AIRE induced antigen expression and antigen presentation in the thymus is a highly tuned system that may be disrupted either by an increased number of different antigens to present, or by increasing the number of the same antigens such that the whole repertoire of antigens is not presented.

Traditionally, the leading view has been that AIRE preferentially induces genes that are tissue specific and that are characterized by low levels of initial expression. ChipSeq studies and other molecular approaches have failed to identify any consensus binding motif in the target DNA. In order to achieve so-called ectopic gene expression, it has therefore been assumed that AIRE interacts with repressive histone marks or repressive chromatin complexes to pinpoint TRAs. Indeed, previous studies have indicated that the majority, but not all, AIRE induced genes have promoters with low levels of H4K3me3 and H3 acetylation [63], but high levels of H3K27me3 [2]. More recently, increased understanding of how AIRE induces its target genes have added even more layers of complexity. In mTECs and transfected cell-lines, AIRE may be located at up to 40 000 sites across the genome at transcriptional start sites, superenhancers and introns [8]. Here, AIRE appear to act rather late in the transcription cycle by reinforcing the actions of other transcription factors and by triggering transcriptional elongation by RNA polymerase [22]. The steps prior to transcriptional elongation, such as chromatin remodulation and accessibility, seem to be AIRE independent [15]. Hence, the AIRE target genes appear to be poised for transcription prior to AIRE induction. Recent evidence suggests that this is achieved in an ordered stochastic manner in which AIRE preferentially targets super enhancers enriched with acetylated H3K27 residues and genomic regions enriched with Z-DNA and/or double-stranded DNA breaks [53, 24, 64]. We consistently find that AIRE induced expression is affecting a large number of genes with little to no initial expression by a large relative increase, but still with a low absolute increase in transcription. However, AIRE seems to also induce an increase in expression on a number of already expressed genes, and while 2/3’s of the genes induced by AIRE WT is categorised as some form of tissue restriction by the Human Protein Atlas, 1/3 is not categorised as such. It is possible the overabundance of AIRE in our transfected system is responsible for these effects, however it may also be secondary expression caused by AIRE induced transcription factors, or that the increase in power of our study allows us to identify effects that may be lost in studies with a lower power focusing on the relative expression of AIRE induced genes.

In summary, we have developed a cell-culture system that allows for in depth analysis of differential transcriptional effects of AIRE variants associated with different clinical phenotypes. By combining a large number of replicates and high-depth RNAseq, we have the statistical power to detect both dramatic and subtle transcriptional differences induced by AIRE WT and how this induction is disrupted in the activity of the pathogenic variants p.R257* and p.C311Y, as well as the increased activity of the common variant p.R471C linked to common organ specific autoimmunity. Consistent with previous studies, we find that AIRE induces thousands of genes from a level of little or no initial expression by a small increase in expression. However, we also find that AIRE is able to boost expression of some already expressed genes, and while the majority of AIRE expressed genes can be categorised as tissue specific, many genes are not. Finally, as a first, we confirm the AIRE regulated expression of a comprehensive number of clinically relevant autoantigens.

## Data availability

The RNAseq data underlying this publication (fastq and bam files) has been submitted to the European Nucleotide Archive with accession number PRJEB83008. RNAseq differential expression results in tables are collected as an Excel file in **Supplementary file B1**. Additional data is available at request.

## Supporting information

Supplemental file B1 RNAseq results

## Acknowledgements

We would like to thank the Research and Development group at the Medical Genetics department of Haukeland University hospital and the Genomics Core Facility at the University of Bergen for help in library preparation and sequencing for RNAseq in this project. This work was supported by grants (to SJ) Helse Vest’s Open Research Grant (grants #912250 and F-12144), the Novo Nordisk Foundation (grant NNF19OC0057445), Research Council of Norway (grant #315599), and (to AHB) an L. Meltzer Foundation project grant. We would also like to thank Haukeland University Hospital, and the University of Bergen for general funding of this project.

